# Rapid evolution in managed grasslands: different land-use regimes are associated with contrasting phenotypes in *Plantago lanceolata*

**DOI:** 10.1101/2020.02.27.967448

**Authors:** Bence Gáspár, Walter Durka, Oliver Bossdorf

**Affiliations:** Plant Evolutionary Ecology, Institute of Evolution & Ecology, University of Tübingen, Auf der Morgenstelle 5, 72076 Tübingen, Germany; Department of Community Ecology, Helmholtz Centre for Environmental Research - UFZ, Theodor-Lieser-Str. 4, 06120 Halle, Germany

**Author notes:** Author for correspondence: Bence Gáspár.

## Abstract

Land-use intensification is a major driver of biodiversity declines, and it is known to decrease species numbers and alter community composition of managed grasslands. An open question is whether similar impacts occur within species, i.e. whether grassland management affects also the genetic diversity of plant populations and alters their genetic composition, possibly reflecting adaptation to land use. To address these questions, we sampled 61 populations of the common grassland herb *Plantago lanceolata* that covered a broad range of land use intensities in the German Biodiversity Exploratories, and we grew their offspring in a common environment to quantify variation in plant size, architecture, reproduction, and leaf economy. All measured traits harboured substantial heritable variation, and six out of nine traits showed population differentiation. Interestingly, several traits were significantly correlated with land use intensity, but with opposing trends for mowing versus grazing: Increased mowing intensity was associated with larger plant size and lower specific leaf area (SLA), which may reflect evolutionary responses to increased light competition and a lesser need for resource conservation in highly productive meadows. In contrast, increased grazing intensity tended to be associated with smaller plant size and higher SLA, a phenotype syndrome known from grazing lawns. Moreover, we found that land-use intensification also affected genetic diversity, again with opposing effects for mowing versus grazing: while increased mowing was associated with decreased levels of intrapopulation phenotypic variation, the opposite was true for increased grazing intensity. In summary, land use intensification has not only already caused rapid evolutionary changes in these grassland populations, it also affects their future evolutionary potential.

## Introduction

Human activity is radically changing the global environment, and it is causing substantial biodiversity declines in all biomes. Recent estimates suggest that species richness has globally decreased by an average of 13.6% since the 16th century, with the greatest declines in the 19th and 20th centuries, and a predicted further 3.4% loss until the end of this century (Newbold *et al*. 2015). Among the many factors contributing to these biodiversity losses, the conversion of natural ecosystems to managed land and the intensification of existing land use practices are among the strongest ones (Sala *et al*. 2000; Foley *et al*. 2005).

While natural biodiversity is impacted by humans, the mechanisms of evolutionary change never cease, new diversity continuously emerges through mutation, and existing diversity is modulated through genetic drift, migration and natural selection. Plant species with broad environmental ranges can be under divergent selection in different environments, and indeed, local adaptation has been widely documented, especially in species with larger populations (Leimu and Fischer 2008). These contrasting environments need not necessarily be located far from each other; in heterogeneous environments divergent selection and local adaptation can take place at very small scales (Linhart and Grant 1996), as it has been shown for example in heavy metal-contaminated soils (Antonovics 1971; Jiménez-Ambriz *et al*. 2007), or even for small-scale herbivore abundances (Dirzo and Harper 1982). Another type of biotic interaction that can create such heterogeneous mosaics are land-use practices in anthropogenic agricultural landscapes, which might strongly differ between neighbouring pieces of land.

Subsistence pastoral systems and the grazing of grasslands by livestock dates back to over 11.000 years, while mowing of meadows for haymaking – a prerequisite for feeding domesticated animals year-round – can be traced back to around 1000 BC (Briggs 2009). Both land-use practices strongly affect the composition of plant communities, however in different ways. Mowing generally creates a uniform, low-canopy, high-light environment, while grazing is heterogeneous on various scales through the selective removal of biomass, and by trampling and depositing dung and urine in patches (Gibson 2009). Researchers repeatedly documented evolutionary responses to different grassland management regimes, starting as early as the 1920’s (Stapledon 1928), in classical work from English grasslands and the Park Grass Experiment (e.g. Warwick and Briggs 1979; Davies and Snaydon 1976; Silvertown *et al*. 2006), and in more recent and contemporary studies (e.g. Suzuki 2008). However, most previous studies included few populations and coarse categories of land-use types, with some exceptions such as two recent studies from the German Biodiversity Exploratories (Völler *et al*. 2013; 2017).

The Biodiversity Exploratories (www.biodiversity-exploratories.de) are a long-term research platform designed for studying the relationships between land use, biodiversity and ecosystem functioning. They provide networks of large numbers of standardised grassland and forest plots across three regions in the north, middle and south of Germany (Fischer *et al*. 2010). In the grassland plots, the intensities of mowing, fertilization and grazing are precisely documented for each plot through annual inventories, providing unique opportunities for studying many populations of the same species along broad gradients of land-use intensities (Blüthgen *et al*. 2012).

Previous studies of land use effects on species diversity often found that species richness of plants decreases with increasing land use intensity, as less competitive species are excluded by dominant competitors at higher nutrient availability (Blüthgen *et al*. 2012). It is still unclear, however, whether similar processes take place at the intraspecific level (Vellend and Geber 2005). In the Biodiversity Exploratories, intraspecific trait variation of grassland plants has been recently studied through a phytometer approach (Herz *et al*. 2017), where multiple species were planted into different plots to demonstrate that changes in land use intensity can create substantial variation in phenotypic traits. However, while this study controlled for plant genotypes to estimate only the plastic components of intraspecific variation, the opposite approach is required if we want to understand adaptation and evolutionary potential: a common-garden approach where genotypes from different origins are planted under controlled environmental conditions, so that only the heritable component of intraspecific variation is remaining. This is the approach we chose for our study with *Plantago lanceolata*.

We chose to work with *Plantago lanceolata*, because it is one of the most widespread grassland species in the Biodiversity Exploratories. Although there has been previous work on *P. lanceolata* in the context of intraspecific variation and land use (Kuiper and Bos 1992), this work was usually restricted to few populations and coarse comparisons of land use types (Warwick and Briggs 1979; van Groenendael 1986; Wolff and Van Delden 1987; van Tienderen and van der Toorn 1991). Here, we studied heritable phenotypic variation in 61 grassland populations of *P. lanceolata* from a broad range of (precisely known) intensities of mowing, fertilization and grazing. We asked the following questions: (i) Do the studied populations harbour significant heritable intraspecific variation? (ii) If yes, how are the mean phenotypes related to land use intensity? (iii) Does land-use intensification also affect the within-population diversity of phenotypic traits?

## Materials and Methods

### Plant material and common-garden experiment

*Plantago lanceolata L*. (Plantaginaceae) is a very common Eurasian grassland rosette herb, distributed over broad geographic and environmental gradients. It is also among the commonest plant species in the Biodiversity Exploratories, occurring on 70% of the grassland plots. The sampling design and experimental setup are depicted and described in detail in Gáspár *et al*. (2018). Briefly, in September 2015, we collected ripe seeds from altogether 61 plots across the three regions of the Biodiversity Exploratories. We used these seeds to establish a common-garden experiment in a greenhouse at the University of Tübingen. The experiment had a randomized block design with three seedlings from each of four mother plants (= four maternal seed families) per sampled plot (total *N* = 741). It was maintained under a 16 h/8 h day/night cycle at 21°C/15°C. After 10 weeks, we harvested all plants, scanned five leaves per plant for further analyses, and dried all plant material at 70°C for at least 72 h. To quantify intraspecific variation in plant size, architecture, reproduction and leaf economy, we measured the following phenotypic traits: (1) aboveground biomass, (2) length of the longest leaf, (3) plant height, (4) the shape (width:length ratio) of leaves and (5) their specific leaf area (SLA; area of five leaves divided by their dry weight), (6) the ‘growth habit’ of the plants (rosette height:width ratio), (7) onset of flowering, (8) number of inflorescences, and (9) reproductive allocation (reproductive:total aboveground biomass ratio).

### Data analysis

We used the R statistical environment (R Development Core Team 2008) for all statistical analyses. Prior to the main analyses, we z-transformed (standardised) all nine response variables by subtracting the mean and dividing by the standard deviation. To simplify our subsequent analyses, and to account for all differences between regions that were of no interest to our study (e.g. altitude, latitude, etc.), we then fitted a linear model for each variable with regions of origin and experimental blocks as main effects, and we used the residuals from these models in all analyses described below (Manning *et al*. 2015; Soliveres *et al*. 2016). To explore overall phenotypic similarities and visualize our data, we ran a principal component analyses (PCAs) on the maternal seed family-level aggregated data using the *vegan* package (Oksanen *et al*. 2017). Next, we tested for heritable variation in phenotypes with two approaches: First, we used the *lme4* package (Bates *et al*. 2015) to fit mixed-effects models that included populations of origin as fixed effect and maternal seed families as random effect. In these models, any significant effect indicates resemblance among relatives and thus heritable variation in a trait. In a second approach, we calculated narrow-sense heritabilities for each trait and population as *h*^2^ = *V*_A_ / (*V*_A_ + *V*_ε_) = 4*V*_FAM_ / (4*V*_FAM_ + *V*_ε_), where *V*_A_ is the additive genetic variance that is equal to four times the variance among families (*V*_FAM_) in half-sib experimental setups, and *V*_ε_ is the residual variance (Petit *et al*. 2001). However, the obtained *h*^2^ values showed a zero-inflated distribution (data not shown), probably because the data were based on only three individuals per family and four families per population – an inevitable consequence of our aim to maximise the number of populations while keeping the total number of samples at a manageable level. As an alternative, we also calculated the phenotypic diversity for each trait and population as the standard deviation of the family-level means within a population. *h*^2^ and phenotypic diversity turned out to be strongly correlated in all traits (*r* = 0.323 to 0.751, all *P* < 0.001), and we therefore used the phenotypic diversity in all further analyses. Finally, to explore relationships between land use intensity and phenotypic traits, we added the vectors of mowing, fertilisation and grazing to the PCAs with the *envfit* function in the *vegan* package, and we related population-level trait means (first aggregated at the family level) or phenotypic diversities, respectively, to mowing, fertilisation and grazing intensity by running separate linear models for each trait and land use component. We corrected all *P*-values for false discovery rates (FDR). The land-use intensity data comes from yearly inventories in the Biodiversity Exploratories, and is based on the number of mowing events, the amount of nitrogen added per hectare, and the number of grazing animals (livestock units) per hectare in a year (for more details see Blüthgen *et al*. 2012).

## Results

The mixed-effect models showed significant population differentiation in six out of the nine studied phenotypic traits, and there were significant maternal seed family effects - indicating heritable variation within populations - in all of the measured traits (Table 1). Across traits, an average of 6.4% of the variance (range 2-11%) resided among populations, and 14.6% (range 11-26%) among families within populations. The first two axes of the principal component analysis explained 55.3% of the multi-trait phenotypic variance and were associated with different traits (Figure 1 and Table 1). Moreover, the PCA indicated three groups of closely related traits: (1) the reproduction-related traits of flowering time, number of inflorescences and reproductive allocation, (2) aboveground biomass and SLA, which were negatively correlated, and (3) the remaining traits, all related to plant architecture and leaf shape.

**Table 1.** Results of mixed-effect models testing for heritable variation among and within 61 natural populations of *Plantago lanceolata*, for phenotypic traits measured in a common environment. The population effects are estimated using family as a random factor, and the family effects are based on four seed families per population. *P*-values are in bold if *P*<0.05 after FDR-correction. The loadings of the first two axes of a principal component analysis are also shown.

|  | Population |  | Seed family | Variance Components |  | PCA loadings |  |
| --- | --- | --- | --- | --- | --- | --- | --- |
|  | *F*-ratio | *P*-value | *P*-value | Among populations | Among families | PC1 (34.9%) | PC2 (20.4%) |
| Leaf shape | 1.164 | 0.221 | **0.008** | 0.02 | 0.11 | -0.38 | -0.11 |
| Leaf length | 1.662 | **0.005** | **0.003** | 0.07 | 0.12 | -0.54 | 0.05 |
| Aboveground biomass | 2.082 | **0.000** | **0.001** | 0.11 | 0.13 | -0.24 | 0.12 |
| # Inflorescences | 1.671 | **0.005** | **0.008** | 0.06 | 0.10 | 0.17 | -0.43 |
| Onset of flowering | 1.658 | **0.006** | **0.000** | 0.09 | 0.26 | 0.06 | 0.56 |
| Reproductive allocation | 1.154 | 0.236 | **0.001** | 0.02 | 0.14 | -0.03 | -0.55 |
| Specific leaf area | 2.152 | **0.000** | **0.003** | 0.11 | 0.12 | 0.29 | -0.35 |
| Plant height | 1.545 | **0.016** | **0.000** | 0.06 | 0.19 | -0.48 | -0.24 |
| Growth habit | 1.336 | 0.076 | **0.000** | 0.04 | 0.14 | -0.39 | -0.05 |

**Figure 1.**
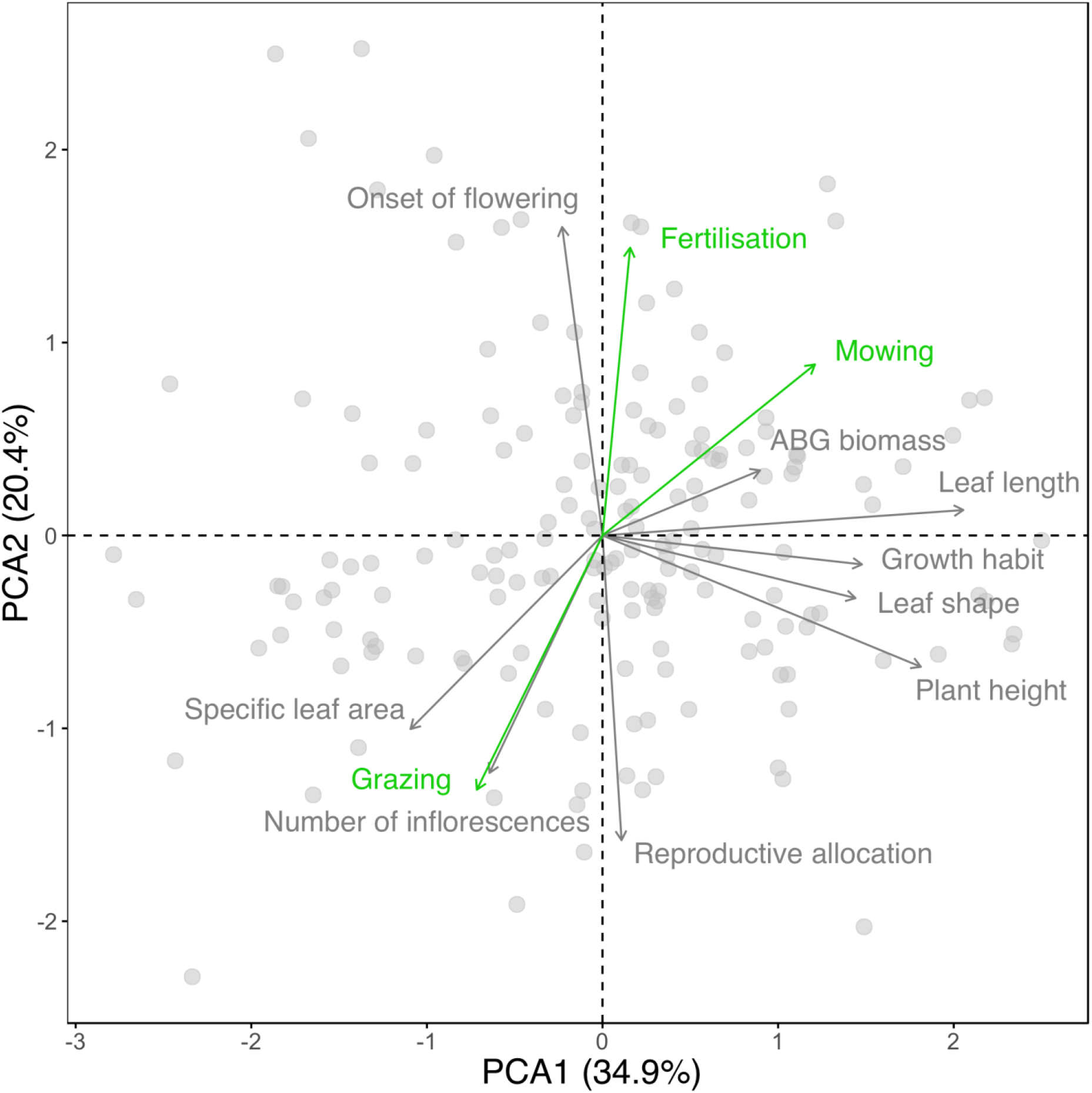
PCA biplot based on maternal seed family-level means of nine phenotypic traits measured in *Plantago lanceolata* in a common environment. Three land-use component vectors are fitted to show correlations between these and the phenotypes.

When we related population-level phenotypic variation to land use intensity, we found the strongest patterns for mowing intensity: at higher mowing intensities, plants were taller, had longer leaves and a greater biomass, but a lower specific leaf area (Table 2). There were also positive relationships between fertilization and biomass, and between grazing intensity and specific leaf area, but these did not remain statistically significant after correcting for false discovery rates (Table 2). An intriguing general pattern was that the signs of relationships tended to be opposite for grazing versus mowing and fertilization (Table 2). For instance, high population-level values of specific leaf area were associated with low levels of mowing intensity but high levels of grazing intensity (Fig. 2). The pattern was further supported by the PCA, where the vectors for mowing and fertilization versus grazing had opposite directions (Fig. 1), i.e. they were associated with contrasting multi-trait phenotypes.

**Table 2.**
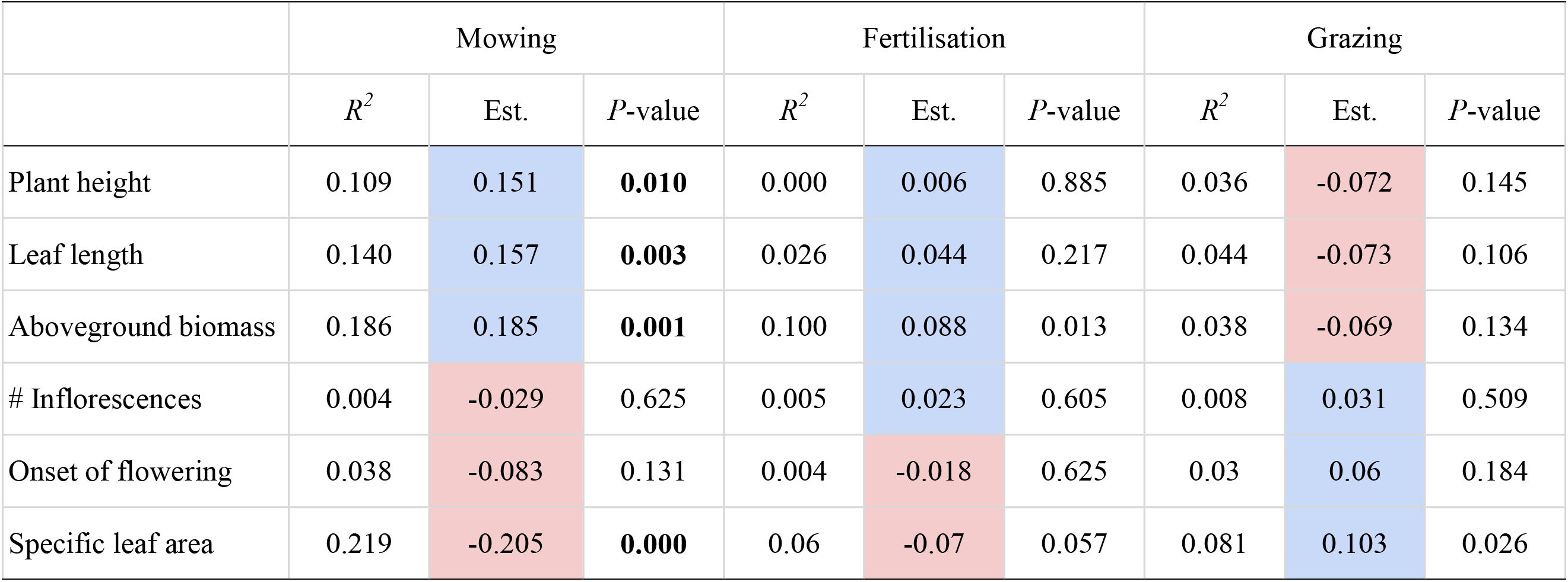
Summaries of linear models testing the effects of mowing, fertilisation or grazing intensities on the population means of six phenotypic traits harbouring heritable variation in *Plantago lanceolata*. *P*-values are in bold if *P*<0.05 after FDR-correction. To show the contrasting effects of mowing/fertilisation vs. grazing, positive regression slopes are highlighted in blue, and negative ones in red.

**Figure 2.**
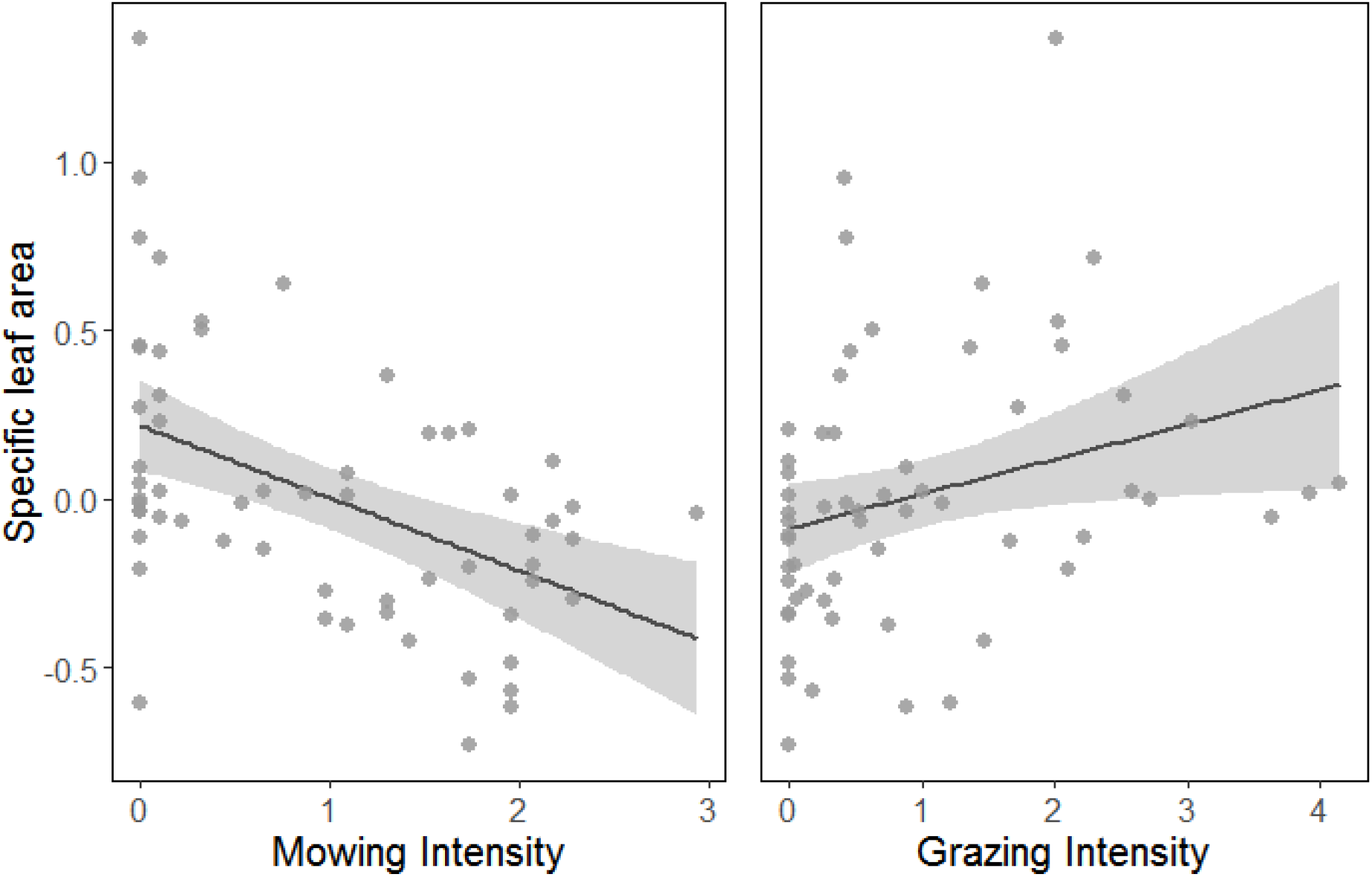
Relationships between land use intensities and average specific leaf area across 60 grassland populations of *Plantago lanceolata*. Mowing and grazing intensities are calculated from the yearly frequency of mowing and grazing animals per hectare, respectively. The fitted generalized linear models (GLMs) and 95% confidence intervals are shown as solid lines and grey shading, respectively.

There were also several patterns of relationships between land use intensities and within-population trait diversities, particularly for mowing intensity (Table 3). Although none of these remained statistically significant after FDR correction, there was again an overall pattern that effects were in opposite directions for grazing versus mowing and fertilization. While higher intensities of mowing and fertilization were generally associated with lower phenotypic diversity, the opposite was true for increasing grazing intensity (Table 3, Fig. 3).

**Table 3.** Summaries of linear models testing the effects of mowing, fertilisation or grazing intensities on population-level phenotypic diversities of nine traits in *Plantago lanceolata*. None of the results were significant after FDR-correction of the *P*-values. To show the contrasting effects of mowing/fertilisation vs. grazing, positive regression slopes are highlighted in blue, and negative ones in red.

|  | Mowing |  |  | Fertilisation |  |  | Grazing |  |  |
| --- | --- | --- | --- | --- | --- | --- | --- | --- | --- |
| | $R^2$ | Est. | *P*-value | $R^2$ | Est. | *P*-value | $R^2$ | Est. | *P*-value |
| Leaf shape | 0.016 | 0.036 | 0.337 | 0.008 | 0.017 | 0.498 | 0.001 | -0.007 | 0.834 |
| Leaf length | 0.002 | 0.014 | 0.713 | 0.002 | 0.008 | 0.733 | 0.039 | -0.048 | 0.125 |
| Aboveground biomass | 0.036 | -0.052 | 0.145 | 0.074 | -0.049 | 0.034 | 0.003 | -0.012 | 0.685 |
| # Inflorescences | 0.100 | -0.099 | 0.013 | 0.058 | -0.049 | 0.061 | 0.045 | 0.055 | 0.102 |
| Onset of flowering | 0.072 | -0.088 | 0.037 | 0.032 | -0.038 | 0.166 | 0.089 | 0.081 | 0.02 |
| Reproductive allocation | 0.079 | -0.091 | 0.028 | 0.027 | -0.035 | 0.202 | 0.061 | 0.066 | 0.054 |
| Specific leaf area | 0.149 | -0.101 | 0.002 | 0.071 | -0.046 | 0.038 | 0.064 | 0.055 | 0.049 |
| Plant height | 0.015 | -0.048 | 0.348 | 0.005 | -0.017 | 0.606 | 0.015 | 0.041 | 0.341 |
| Growth habit | 0.003 | -0.014 | 0.701 | 0.013 | 0.02 | 0.379 | 0.002 | 0.009 | 0.748 |

**Figure 3.**
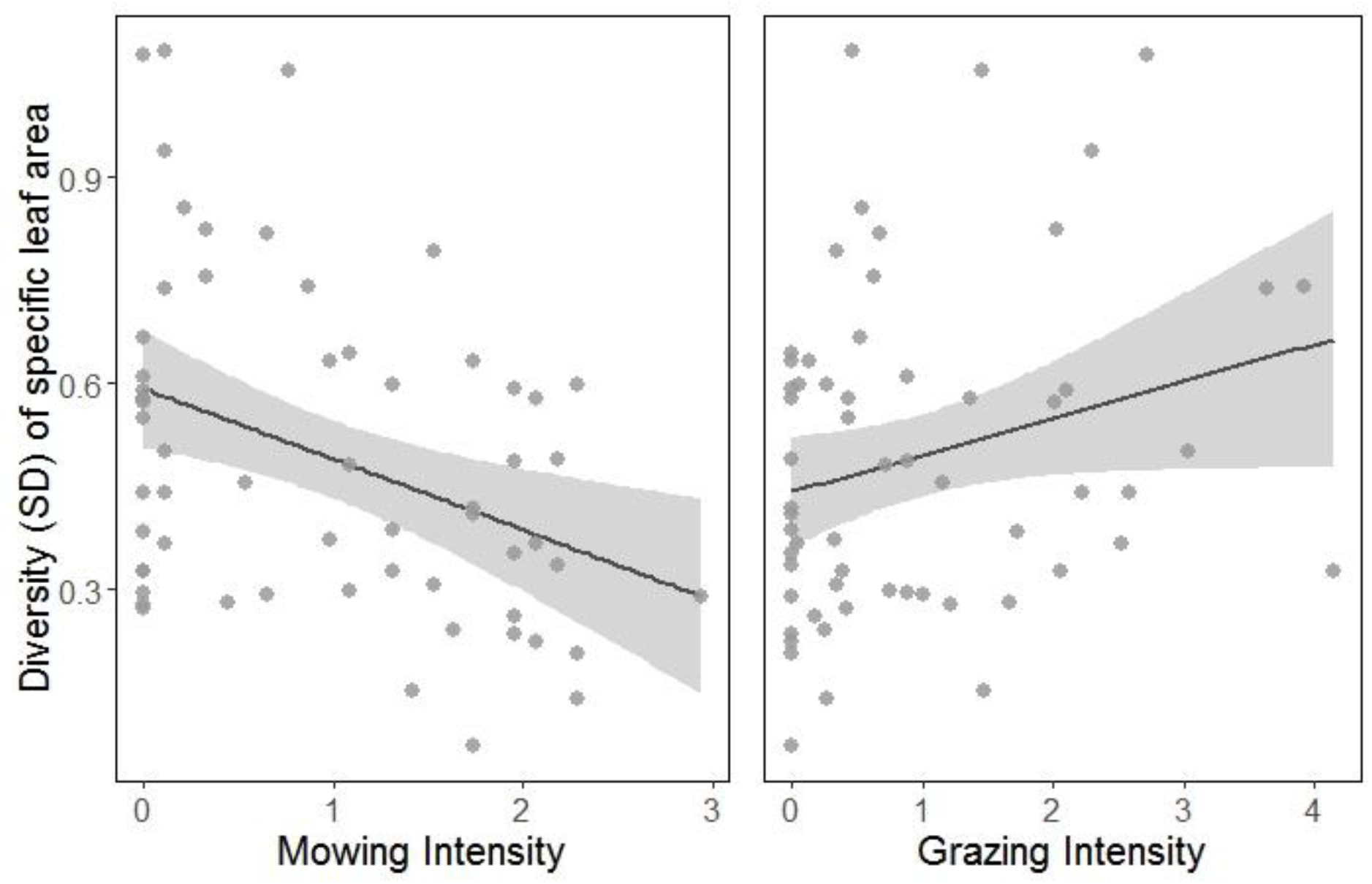
Relationships between land use intensities and the within-population diversities of specific leaf area across 60 grassland populations of *Plantago lanceolata*. Mowing and grazing intensities are calculated from the yearly frequency of mowing and grazing animals per hectare, respectively. The fitted generalized linear models (GLMs) and 95% confidence intervals are shown as solid lines and grey shading, respectively.

## Discussion

Land use intensification and its ecological and evolutionary consequences are important research topics in current evolutionary ecology. Here, we investigated whether the common grassland plant *Plantago lanceolata* is undergoing evolutionary changes in response to land-use intensification. We found substantial heritable phenotypic variation - indicating evolutionary potential - in all studied traits, and that there were some significant associations between phenotypes and land use intensity. Most interestingly, throughout our analyses, mowing and fertilization tended to be associated with contrasting phenotypes, and they also had opposing effects on within-population phenotypic diversity, suggesting that these two land use processes have fundamentally different effects on the evolution of these plant populations.

Several previous studies already studied intraspecific variation of *P. lanceolata* (e.g. Sagar and Harper 1964; Warwick and Briggs 1979; Kuiper and Bos 1992), or relationships between intraspecific variation and environment of origin. One classic study showed substantial population differentiation as well as within-population variation in five reproductive traits among eight natural *P. lanceolata* populations (Primack and Antonovics 1981). Other studies found strong heritable differences between plants from two closely located contrasting habitats (van Groenendael 1986), and even local adaptation with consistent better survival at the home site (Van Tienderen and van der Toorn 1991). Another previous study found substantially higher population differentiation in four of the traits we also measured (Wolff and Van Delden 1987). However, some of this discrepancy may result from differences in experimental design, since Wolff and Van Delden (1987) studied F_2_ plants from full-sib families (from only two pastures and two meadows), whereas we worked with F_1_ individuals from half-sib families that – as a result of wind pollination and obligate outcrossing – still contained random paternal alleles, and therefore harboured greater within-population variance. In summary, while estimates of phenotypic variation tended to be lower in our study than in some previous studies on *P. lanceolata*, our results confirmed the existence of significant, and potentially adaptive, heritable variation in phenotype.

### Land use and population-level phenotypic trait means

When examining patterns of phenotypic variation in relation to land use intensity, the most intriguing result were the contrasting effects with regard to grazing and mowing. Plants originating from sites with high mowing intensity were in general larger (in terms of aboveground biomass, length of the longest leaf and height of the longest inflorescence stalk) and they had a lower specific leaf area, whereas plants from sites with high grazing intensity tended to be smaller, with higher SLA. These opposite effects of the two land use types consistently appeared across almost all of the measured traits. Trait relationships with fertilisation intensity followed mostly the same direction as mowing; it is well known from the Biodiversity Exploratories that these two land-use components are strongly correlated (Blüthgen *et al*. 2012). There are some previous studies of *P. lanceolata* comparing different land use types (van Groenendael 1986; Wolff and Van Delden 1989; Kuiper and Bos 1992) which also found that a prostrate growth habit with more dormant seeds is associated with grazing, whereas in meadows there are taller and larger plants with more, readily germinating seeds. However, these studies compared only a handful of sites with discrete categories of land use, often confounded with other environmental factors (e.g. grazed sand dunes vs. wet, nutrient-rich hayfields). Our study corroborates these findings across a much larger number of natural populations and broad gradients of mowing and grazing intensities. Similar to ours, other studies from the Biodiversity Exploratories showed that patterns of phenotypic trait differentiation were strongest in relation to mowing (Völler *et al*. 2013; Völler *et al*. 2017).

We found that lower values of specific leaf area were associated with larger plants and more intense mowing, but that SLA increased with decreasing plant size and increasing grazing. At first glance this result is surprising because higher SLA is in generally thought to be associated with higher relative growth rate, leaf nitrogen content and shading (Lambers and Poorter 1992). However, most of these relationships have been observed across species, often with a strong emphasis on woody species (Wright *et al*. 2004; Poorter *et al*. 2009; Díaz *et al*. 2016). Several previous studies quantified intraspecific variation in SLA but only the plastic components of it, or were unable to separate plastic and heritable components, e.g. where intraspecific variation was measured in different habitats in the field (e.g. Shipley and Almeida-Cortez 2003; Bilton *et al*. 2010; Hulshof *et al*. 2013; Jung *et al*. 2014). We are aware of only one other study explicitly focusing on heritable intraspecific variation in SLA in a grassland plant: Scheepens *et al*. (2010) conducted a reciprocal transplant experiment with *Campanula thyrsoides* across different altitudes, and they found that there was significant heritable differentiation in SLA among populations.

Some previous studies might help to explain the observed relationships between SLA, plant size and land use. A multi-species study across 157 species showed that in the majority of these, including 14 out of 20 studied herbaceous species, SLA decreased with increasing leaf area (Milla and Reich 2007). The authors explained this by the increased need for structural support in larger plants. In the Biodiversity Exploratories, more intensely mown plots are usually also more fertilised (Blüthgen *et al*. 2012), with more intense competition for light (Kuiper and Bos 1992, page 273), resulting in larger and taller plants that must invest more in structural support, thereby decreasing SLA. On the mown plots, nutrients are readily supplied by fertilisation, so there is no selection for resource conservation, but the race for light - following each mowing that “cleans the slate” homogeneously across the whole plot - might select for taller plants with bigger and stronger leaves. Another reason why SLA of *P. lanceolata* is positively related to grazing could be that the trait is functionally related to herbivory. Plants with high SLA are often not only shorter-lived but also more palatable (Reich *et al*. 1999; Poorter *et al*. 2009). If these plants have a high tolerance of grazing, i.e. they regrow better than competitors or even overcompensate, then grazing will select for high SLA. In a multispecies study from southern Patagonia, Cingolani *et al*. (2005) indeed showed that a positive feedback loop can increase the abundance of preferentially eaten plants. They argued that this generally requires three conditions to be met: high levels of grazing tolerance, herbivore preference, and high resource availability. *P. lanceolata* is known to be grazing-tolerant but preferentially grazed by cattle and sheep (Sagar and Harper 1964). It is therefore possible that this explanation is relevant for our study system, and grazing selects for small but rapidly regrowing genotypes with high SLA.

We did not find land-use related population differentiation in growth habit (related to leaf angle), as it was the case in earlier studies of *Plantago lanceolata* (van Groenendael 1986; Wolff and Van Delden 1989). One explanation for this could be that our F_1_ plant material with random fathers had too high levels of within-family variability, compared to several generations of selective breeding in a previous experiment that found strong heritability of this trait (Wolff and Van Delden 1989). Another explanation might be that, unlike previous studies with simple comparisons of land-use categories, we analysed gradients of land-use intensity, and relationships between leaf angle and land use might be complex or non-linear.

### Genetic diversity in phenotypic traits

All phenotypic traits analysed in our study showed significant variation at the level of seed families within populations, which allowed us to examine relationships between within-population phenotypic diversity and land use. We found that mowing generally decreased phenotypic diversity, while grazing tended to increase it. The most likely explanation for these contrasting results is that mowing and fertilization generally homogenise environmental conditions across a meadow, whereas grazing creates heterogeneity and a greater diversity of microhabitats (Bakker *et al*. 1984). Although the extent of the heterogenising effect of grazing depends on the quality of grazing (e.g. livestock density and selective browsing of different livestock species), the spatial structure of vegetation, and the scale of study (Adler *et al*. 2001), pastures as the ones studied here typically experience more heterogeneous biomass removal, nutrient supply, disturbance, and competition than similar, but mown, grasslands (Bakker *et al*. 1983; Bakker *et al*. 1984; McNaughton 1984). Further evidence for this comes from a recent study in the Biodiversity Exploratories that found land use, especially mowing intensification, to cause biotic homogenisation across 12 trophic groups (Gossner *et al*. 2016). In summary, we found that increased mowing decreased within-population phenotypic diversity in *Plantago lanceolata* populations, but that the opposite was true for increased grazing, and we suggest that these contrasting results are explained by the heterogenising qualities of grazing versus homogenising effects of mowing and fertilisation.

## Conclusions

Our study demonstrates that increased mowing and grazing affects evolution and diversity of phenotypes in a common grassland plant. On the one hand, different land use practices were associated with contrasting multi-trait phenotypes, most likely reflecting adaptation to the selection regimes exerted by these land use practices. On the other hand, mowing and grazing had opposing effects on within-population diversity, presumably because mowing makes habitat conditions more homogenous whereas the opposite is true for grazing. Taken together, land use intensification not only causes rapid evolution of phenotypes in the studied grassland populations, but it also affects their future evolutionary potential.

## Authors’ contributions

OB and WD planned and designed the research; BG conducted fieldwork and performed experiments; BG and OB analysed the data and wrote the manuscript.

## Acknowledgements

This work has been funded through the DFG Priority Program 1374 ‘Biodiversity Exploratories’ (DFG grants DU 404/9-1 and BO 3241/2-1). We thank the managers of the three Exploratories, Kirsten Reichel-Jung, Katrin Lorenzen and Martin Gorke, and all former managers, for their work in maintaining the plot and project infrastructure, Christiane Fischer and Jule Mangels for their support through the central office, Andreas Ostrowski and Michael Owonibi for managing the central database, and Markus Fischer, Eduard Linsenmair, Dominik Hessenmöller, Daniel Prati, Ingo Schöning, Francois Buscot, Ernst-Detlef Schulze, Wolfgang Weisser and the late Elisabeth Kalko for their role in setting up the Biodiversity Exploratories project. Fieldwork permits were issued by the responsible state environmental offices of Baden-Württemberg, Thüringen, and Brandenburg (according to § 72 BbgNatSchG). We are grateful to Florian Frosch, Jan Helbach for their help during the field campaign, to Christiane Karasch-Wittmann, Eva Schloter, Sabine Silberhorn, and Zhiyong Liao for their technical assistance at the University of Tübingen, and to Madalin Parepa and Niek Scheepens for their support with data analyses.

---

**Table S1.** Standardised land-use information of the study sites calculated from the frequency of mowing (y^−1^), amount of nitrogen added to the plots (kg ha^−1^ y^−1^), and density of grazing animals (livestock unit ha^−1^ y^−1^), averaged across the years 2006–2014.

| Schwäbische Alb |  |  |  | Hainich-Dün |  |  |  | Schorfheide-Chorin |  |  |  |
| --- | --- | --- | --- | --- | --- | --- | --- | --- | --- | --- | --- |
| Plot ID | Mowing | Fertilisation | Grazing | Plot ID | Mowing | Fertilisation | Grazing | Plot ID | Mowing | Fertilisation | Grazing |
| AEG02 | 2.85 | 7.56 | 0 | HEG05 | 1.97 | 3.51 | 0.95 | SEG06 | 0.39 | 0 | 1.67 |
| AEG03 | 1.97 | 1.17 | 0.05 | HEG06 | 1.47 | 1.75 | 0.40 | SEG13 | 1.96 | 0.71 | 0.17 |
| AEG12 | 2.16 | 2.43 | 0 | HEG07 | 0.20 | 0.03 | 2.43 | SEG25 | 1.97 | 0 | 0 |
| AEG13 | 2.06 | 2.54 | 0 | HEG09 | 0 | 0 | 0.55 | SEG30 | 1.77 | 0 | 0 |
| AEG14 | 1.97 | 3.71 | 0 | HEG11 | 0.98 | 0.96 | 0.13 | SEG31 | 1.77 | 0 | 0 |
| AEG15 | 2.95 | 5.36 | 0 | HEG14 | 1.28 | 3.40 | 0.40 | SEG32 | 1.77 | 0 | 0 |
| AEG17 | 2.26 | 1.7 | 0 | HEG17 | 0.10 | 0 | 0.41 | SEG33 | 0.33 | 1.08 | 2.03 |
| AEG23 | 1.77 | 0.34 | 0 | HEG18 | 0.11 | 0 | 0.46 | SEG34 | 0.49 | 1.67 | 1.15 |
| AEG25 | 0 | 0.17 | 0.41 | HEG20 | 0 | 0 | 0.87 | SEG36 | 0 | 0 | 2.17 |
| AEG26 | 0 | 0 | 2.01 | HEG26 | 1.18 | 0.63 | 0 | SEG37 | 0.2 | 0 | 2.84 |
| AEG27 | 0 | 0 | 1.2 | HEG27 | 1.28 | 1.91 | 0.04 | SEG38 | 0.88 | 0 | 3.98 |
| AEG29 | 1.08 | 0.45 | 0.73 | HEG29 | 1.57 | 1.16 | 0.25 | SEG39 | 0.59 | 0.2 | 0.76 |
| AEG30 | 0.79 | 0.31 | 1.41 | HEG34 | 1.47 | 2.01 | 0.40 | SEG40 | 0 | 0 | 4.28 |
| AEG31 | 0.69 | 0 | 0.99 | HEG41 | 0 | 0 | 0.87 | SEG41 | 0.1 | 0 | 4.2 |
| AEG33 | 0 | 0 | 1.27 | HEG43 | 0.39 | 0 | 0.61 | SEG44 | 0 | 0.32 | 2.09 |
| AEG35 | 2.16 | 1.34 | 0 | HEG44 | 0.29 | 0 | 0.54 | SEG45 | 0 | 0 | 1.81 |
| AEG38 | 1.97 | 0.24 | 0 | HEG45 | 0 | 0 | 0.47 | SEG47 | 0.1 | 0 | 2.3 |
| AEG39 | 2.06 | 1.90 | 0 | HEG48 | 1.18 | 0.29 | 0.69 | SEG48 | 0.1 | 0 | 2.81 |
| AEG40 | 2.36 | 1.63 | 0.23 | HEG49 | 1.38 | 1.22 | 0.24 | SEG49 | 0 | 0 | 2.61 |
| AEG41 | 2.36 | 4.62 | 0.05 | HEG50 | 0.98 | 1.36 | 0.32 | SEG50 | 0 | 0 | 1.98 |
| AEG42 | 1.47 | 2.10 | 1.34 |  |  |  |  |  |  |  |  |

**Table S2.** Sample sizes in the final analyses (initial N = 741) and summary statistics of the raw data

|  | N |  |  | Trait summary statistics |  |  |  | Variance components |  |  |
| --- | --- | --- | --- | --- | --- | --- | --- | --- | --- | --- |
|  | Individuals | Families | Populations | Mean | SD | Min | Max | EP | Fam | Residual |
| Leaf shape | 731 | 247 | 61 | 9.2 | 2.18 | 4.79 | 16.9 | 0.02 | 0.11 | 0.87 |
| Leaf length | 730 | 247 | 61 | 26.3 | 4.13 | 13 | 44 | 0.07 | 0.12 | 0.82 |
| Aboveground biomass | 731 | 247 | 61 | 8.44 | 2.25 | 0.35 | 14.22 | 0.11 | 0.13 | 0.77 |
| # Inflorescences | 731 | 247 | 61 | 13.39 | 5.58 | 0 | 36 | 0.06 | 0.1 | 0.83 |
| Onset of flowering | 654 | 231 | 61 | 58.87 | 8.29 | 45 | 85 | 0.09 | 0.26 | 0.65 |
| Reproductive allocation | 731 | 247 | 61 | 0.42 | 0.17 | 0 | 0.71 | 0.02 | 0.14 | 0.84 |
| Specific leaf area | 731 | 247 | 61 | 39.44 | 10.83 | 19.07 | 83.82 | 0.11 | 0.12 | 0.78 |
| Plant height | 701 | 241 | 61 | 53.62 | 8.68 | 11 | 75 | 0.06 | 0.19 | 0.75 |
| Growth habit | 731 | 247 | 61 | 0.48 | 0.16 | 0.11 | 1.11 | 0.04 | 0.14 | 0.82 |

